# The time course of categorical and perceptual similarity effects in visual search

**DOI:** 10.1101/2022.05.16.492133

**Authors:** Lu-Chun Yeh, Marius V. Peelen

## Abstract

During visual search for objects (e.g., an apple), the surrounding distractor objects may share perceptual (tennis ball), categorical (banana), or both (peach) properties with the target. Previous studies showed that the perceptual similarity between target and distractor objects influences visual search. Here, we tested whether categorical target-distractor similarity also influences visual search, and how this influence depends on perceptual similarity. By orthogonally manipulating categorical and perceptual target-distractor similarity, we could investigate how and when the two similarities interactively affect search performance and neural correlates of spatial attention (N2pc) using electroencephalography (EEG). Behavioral results showed that categorical target-distractor similarity interacted with perceptual target-distractor similarity, such that the effect of categorical similarity was strongest when target and distractor objects were perceptually similar. EEG results showed that perceptual similarity influenced the early part of the N2pc (200-250 ms after stimulus onset), while categorical similarity influenced the later part (250-300 ms). Mirroring the behavioral results, categorical similarity interacted with perceptual similarity during this later time window, with categorical effects only observed for perceptually similar target-distractor pairs. Together, these results provide evidence for hierarchical processing in visual search: categorical properties influence spatial attention only when perceptual properties are insufficient to guide attention to the target.

**Public significance statement:** Searching for a target object among perceptually similar distractor objects (e.g., looking for an apple among peaches) is relatively difficult. In daily life, target and distractor objects may not only share perceptual but also non-perceptual properties, such as category membership (e.g., fruit). Here, we show that categorical similarity between target and distractor objects influences search performance, particularly when target and distractor objects are perceptually similar. Using electrophysiological recordings, we demonstrate that attentional selection is first influenced by perceptual and then by categorical information. This later categorical influence depended on perceptual similarity, being strongest for perceptually similar objects. These findings provide evidence for hierarchical processing in visual search, with categorical properties extracted after perceptual properties.

## Introduction

Searching for an apple among bananas is easier than searching for an apple among peaches. Behavioral studies have shown that search performance is strongly influenced by the perceptual similarity of target and distractor objects (Alexander & Zelinsky, 2012; Barras & Kerzel, 2017; Duncan & Humphreys, 1989; Proulx & Egeth, 2006; Treisman & Gelade, 1980; Yeh et al., 2019). This effect partly reflects differences in top-down spatial attentional orienting to the target during visual search (Aubin & Jolicoeur, 2016; Eimer, 1996; Luck, 1995; Luck & Hillyard, 1994; Yeh et al., 2019). In those studies, spatial attention is indexed by the N2pc component, a lateralized EEG component over posterior electrodes between 200-300 msec after display onset (Eimer, 1996; Luck & Hillyard, 1994). Like targets, perceptually similar nontargets also elicited an N2pc in displays where the target was absent (Aubin & Jolicoeur, 2016; Luck, 1995; Luck & Hillyard, 1994). Moreover, when the target was presented with perceptual similar distractors in the opposite hemifield, the target-related N2pc was reduced (Yeh et al., 2019). These findings can be explained by theories that propose that top-down search templates bias visual processing and guide attention to possible target locations (Desimone & Duncan, 1995; Duncan & Humphreys, 1989; Wolfe, 2007; Wolfe, 2021). Distractors that share features with the target are more distracting, i.e., pull spatial attention away from the target, because they more closely resemble the search template.

Unlike the stimuli used in most laboratory studies, however, real-world objects also have non-perceptual properties, such as their category (e.g., being animate or inanimate), associated context, and non-observable physical properties (e.g., weight). What role do these properties play in visual search? Interestingly, distractors that share non-perceptual properties with the target have been shown to attract attention and influence search performance (Belke et al., 2008; Guo et al., 2020; Moores et al., 2003; Seidl-Rathkopf et al., 2015; Telling et al., 2010; Wyble et al., 2013). For example, distractor objects that were categorically associated with the target (e.g., table when searching for chair) were more often fixated than non-associated objects (Moores et al., 2003). Furthermore, EEG studies have shown that the N2pc component is modulated by the semantic similarity (including categorical similarity) of the target and distractors (Telling et al., 2010), and that targets defined at the category level (e.g., clothing) elicit an N2pc (Nako et al., 2014; for similar results for alphanumerical categories, see Nako et al., 2016; Nako, Wu, & Eimer, 2014; Wu et al., 2013). Altogether, these results suggest that the semantic category of objects influences visual search and modulates the N2pc, much like perceptual guidance mechanisms.

However, before concluding that abstract object properties guide search similar to perceptual properties, it is important to consider whether categorical influences on search could be mediated by perceptual guidance mechanisms (Wolfe & Horowitz, 2017). For example, target templates may spread to activate perceptual features of target-associated items, e.g., searching for a table may activate the perceptual representation of a chair as well (Moores et al., 2003; Telling et al., 2010). Furthermore, categorical templates may consist of perceptual features that are common to a category (Yang & Zelinsky, 2009) or of a set of multiple distinct category-diagnostic features (Nako et al., 2016; Reeder & Peelen, 2013; Treisman, 2006). Finally, because real-world object categories are often correlated with perceptual features (e.g., animals typically have more curvilinear features than manmade objects), what may appear as a categorical similarity effect may instead reflect residual perceptual similarity differences, with same-category distractors being perceptually more similar to the target than different-category distractors (Alexander & Zelinsky, 2011; Mohan & Arun, 2012). These considerations raise the question of how categorical similarity effects are related to (and may interact with) perceptual similarity effects during visual search.

In the present behavioral and EEG experiments, we investigated the distinct and interactive influence of categorical and perceptual similarity on visual search by manipulating categorical and perceptual similarity in a factorial design. To this aim, we used a stimulus set consisting of animate and inanimate objects that were perceptually similar or dissimilar (Figure 1A). Participants were cued to search for one specific object (e.g., “snake”), which was presented together with one distractor that could be from the same (e.g., snail) or different (e.g., rope) category, and could be perceptually similar (e.g., rope) or dissimilar (e.g., airplane). This design allowed us to test for the independent and interactive effects of perceptual and categorical similarity on visual search performance and lateralized EEG responses.

**Figure 1.**
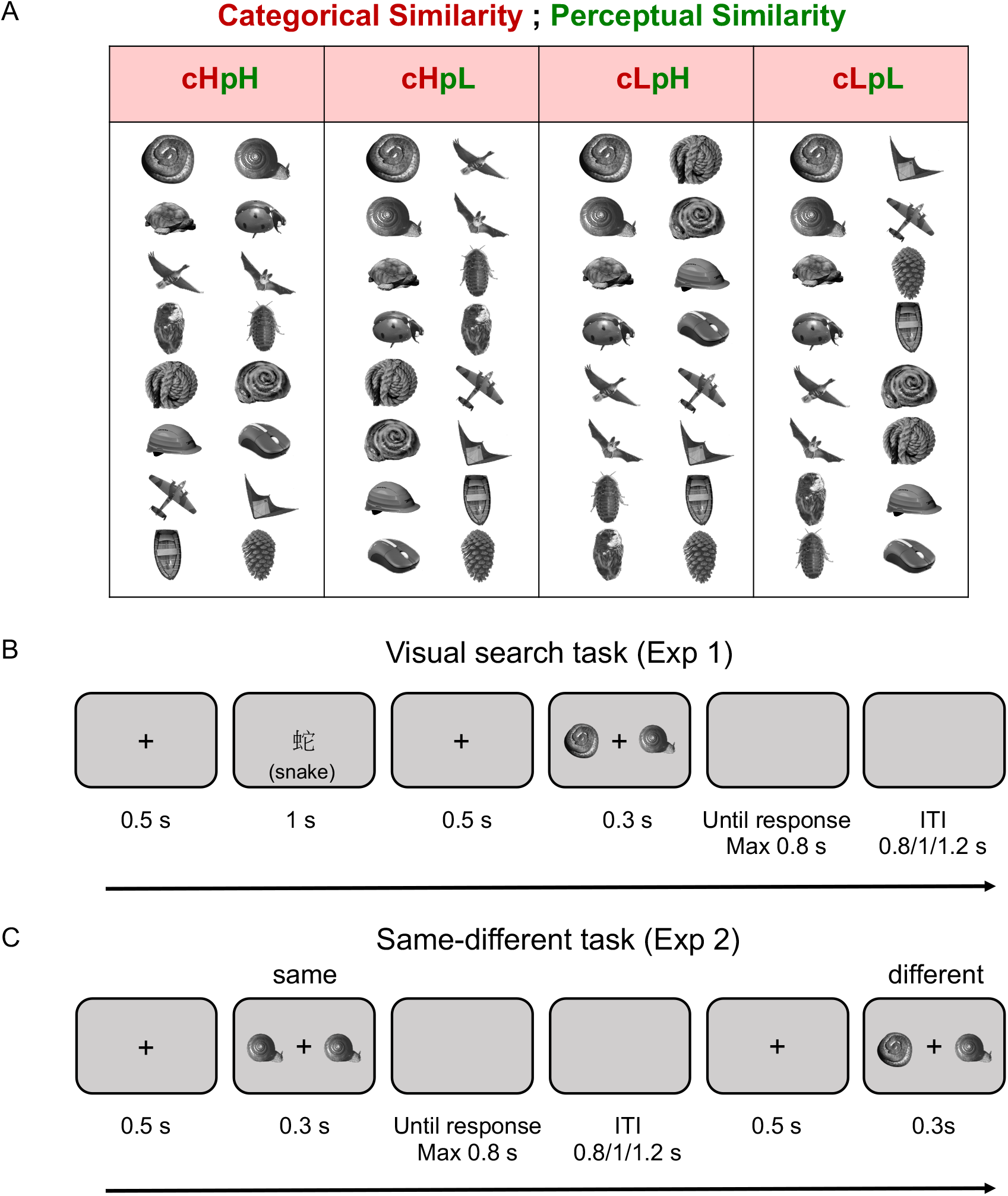
(A) Stimulus pairs for the four search conditions. There are eight pairs for each condition. Four exemplars were used for each object (e.g., four snails); one is shown here. (c: categorical similarity, p: perceptual similarity, H: high similarity, L: low similarity. E.g., cHpH indicates high categorical and high perceptual similarity). (B) Illustration of trial procedure for the visual search task. Participants were instructed to search for a predefined target (in Chinese) from within a search array and to press the mouse button mapping to the target location (ITI = intertrial interval). The cue in English was not shown. (C) Illustration of trial procedure for the same-different task (Experiment 2). Participants were instructed to indicate whether the two objects were the same or different.

A previous fMRI study mapped out the perceptual and categorical representations of the same stimulus set (Proklova et al., 2016). In that study, perceptual object properties were encoded in more posterior visual cortex regions than object categories, indicating hierarchical processing, with perceptual attributes processed before categorical attributes. Accordingly, we predicted that perceptual properties would influence attentional allocation (and the N2pc) before categorical properties would. This led to the key prediction that categorical and perceptual similarity should interact: if the target and distractors are perceptually *dissimilar*, the target can quickly be selected based on its unique perceptual properties, such that the distractor may not be processed up to the category level. By contrast, if the target and distractor are perceptually similar, perceptual guidance is no longer effective, such that categorical properties can influence the search process (i.e., slower search when the distractor is of the same category as the target).

## Experiment 1

### Methods

#### Participants

Experiment 1 was a pilot study for the following EEG study. Sample size was determined by the number of available participants during the testing week. Seventeen healthy volunteers (10 females, mean age = 22.59, SD = 2.99 years) participated in this study. All participants had a normal or corrected-to-normal vision. The participants gave informed written consent and received monetary compensation for their time (160 NTD, about 6 USD). The study was approved by the research ethics office of the National Taiwan University. We excluded one participant’s data from the analyses because his/her overall accuracy was below three standard deviations of the mean.

#### Stimuli

We used sixteen objects identical to the ones used in an earlier fMRI study from our group (Proklova et al., 2016). Half of the objects were animate, and the other half were inanimate. Additionally, there were four exemplars of each object, for example, four images of a snail, resulting in 64 stimuli in total. All stimuli were gray scaled and presented on a gray background. Each stimulus was fitted into an invisible frame subtending a visual angle of approximately 8° (height) × 8° (width). Stimuli were presented using the E-Prime software (Psychology Software Tools, Inc.) and presented on a 17-in. cathode ray tube monitor with a refresh rate of 60 Hz.

#### Task design and procedure

The experimental design followed a 2 (categorical similarity: high, low) × 2 (perceptual similarity: high, low) within-subjects factorial design. There were eight pairs of objects for each condition, resulting in a total of 32 pairs of stimuli (Figure 1A). There were two objects in each pair, one as a target and the other as a distractor. The two objects were both animate or both inanimate for high categorical similarity trials, and they were one animate and one inanimate for low categorical similarity trials. The two objects had a similar shape for high perceptual similarity trials, but not for low perceptual similarity trials.

In the beginning of a trial, a 500-msec fixation cross was presented at the center of the screen to signal the onset of the trial. Then, a target cue (in Chinese) was presented for 1000 msec and followed by another fixation for 500 msec. After that, a search display consisting of two objects was presented for 300 msec and followed by a maximal 800 msec blank for response. Participants were instructed to localize the target indicated by the cue. They responded using both hands by pressing the two mouse buttons with their thumbs, right button for the target on the right and left button for the left. The target locations were fully counterbalanced. The inter-trial intervals varied randomly between 800, 1000, and 1200 msec. The task procedure is illustrated in Figure 1B.

#### General experimental procedure

Participants sat comfortably in a dimly illuminated room, at a distance of 57 cm from the monitor. Then, participants received written and verbal instruction about the task requirements. After that, participants practiced a block of 8 trials showing the eight different objects not used in the formal experiment. During the actual experiment, participants were required to fixate on a cross at the center of the screen and respond as quickly and accurately as they could. Trials were presented in 32 blocks of 32 trials each, yielding a total of 1024 trials (256 trials for each condition). In each block, all 32 pairs (8 pairs for each condition) were included, and each stimulus was defined as a target twice. However, which stimulus was defined as a target and presented on the left or right side were counterbalanced across blocks. The combination of the stimulus pairs using the different exemplars was also counterbalanced across blocks. The trial order within blocks was randomized. Participants could take a rest between blocks and they could self-initiate the next block. The full experimental time for each participant was between 60 to 80 minutes.

#### Behavioral analyses

The behavioral analyses were conducted using JASP (Version 0.16.0.0). All correct trials were included in the RT analyses (no outlier removal). Accuracy and RTs were analyzed by two-way (categorical similarity: high, low x perceptual similarity: high, low) repeated-measures ANOVA.

### Results

The RT data (Figure 2A) revealed significant main effects of both categorical similarity, *F*(1, 15) = 39.556, *p* < .001, *η^2^_p_* = 0.725 and perceptual similarity, *F*(1, 15) = 102.996, *p* < .001, *η^2^_p_* = 0.873. Responses in the low categorical similarity condition (*M* = 451.321 msec, *SE* = 23.661 msec) were faster than in the high categorical similarity condition (*M* = 472.457 msec, *SE* = 25.970 msec), and responses in the low perceptual similarity condition (*M* = 438.561msec, *SE* = 23.491 msec) were faster than in the high perceptual similarity condition (*M* = 485.218 msec, *SE* = 26.218 msec). Importantly, there was a significant two-way interaction between categorical and perceptual similarity, *F*(1, 15) = 5.898, *p* = .028, *η^2^_p_* = 0.282. To follow up on the interaction, we tested the categorical effect separately for low and high perceptual similarity conditions (Figure 2B) using paired sample *t*-tests (with Bonferroni correction for two tests, *p*<.025). The results showed significant categorical effects for both high (*t*(15) = 6.578, *p* < .001, *M*_difference_ = 26.599 msec, *SE* = 4.983 msec) and low perceptual similarity conditions (*t*(15) = 3.876, *p* < .001, *M*_difference_ = 15.674 msec, *SE* = 2.807 msec).

**Figure 2.**
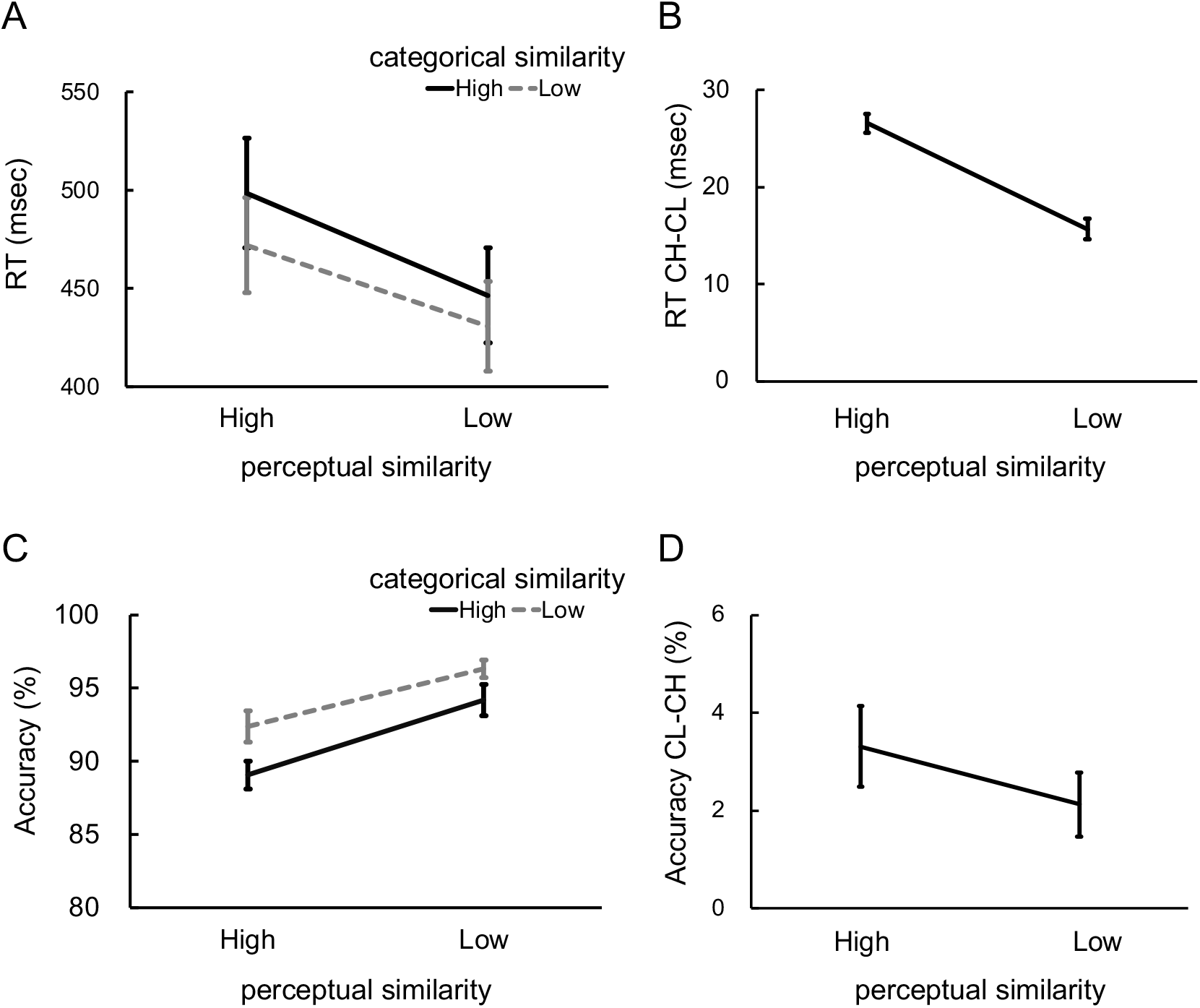
Behavioral results including RT (top panel) and accuracy (bottom panel) from Experiment 1. Error bars represent the standard error of the mean. (A) Mean reaction times for all four conditions. (B) The categorical effect (low-high categorical similarity) of reaction time separately for high and low perceptual similarity conditions. (C) Mean accuracy results for all four conditions. (D) The categorical effect (low-high categorical similarity) of accuracy time separately for high and low perceptual similarity conditions.

The accuracy data (Figure 2C) showed significant main effects of both categorical similarity, *F*(1, 15) = 28.165, *p* < .001, *η^2^_p_* = 0.652, and perceptual similarity, *F*(1, 15) = 30.658, *p* < .001, *η^2^_p_* = 0.671. As shown in Figure 2D, the low categorical similarity condition (*M* = 94.344%, *SE* = 0.747%) was more accurate than the high categorical similarity condition (*M* = 91.625%, *SE* = 0.815%), and the low perceptual similarity condition (*M* = 95.25%, *SE* = 0.757%) was more accurate than the high perceptual similarity condition (*M* = 90.719%, *SE* = 0.923%). There was no significant interaction between categorical and perceptual similarity, *F*(1, 15) = 1.215, *p* = .288, *η^2^_p_* = 0.075.

### Discussion

In Experiment 1, we found that categorical similarity influenced search performance, with increasing similarity decreasing search performance. These results are consistent with previous studies showing that high-level conceptual or semantic information can bias attention in visual search (Belke et al., 2008; de Groot et al., 2016; Moores et al., 2003). In line with these studies, these results support the idea that attention is influenced not only by low-level visual features but also by high-level categorical information (Jenkins et al., 2018; Kaiser et al., 2016; Nako et al., 2016; Nako, Wu, & Eimer, 2014). Importantly, however, we found that the categorical similarity RT effect depended on perceptual similarity, being almost twice as large when the target and distractor objects were perceptually similar. This result is in accordance with our hypothesis that categorical information influences search performance more strongly when perceptual information is less efficient in guiding attention to the target.

However, an alternative explanation for the effect of categorical similarity could be that the objects in the high and low categorical similarity conditions were not equally discriminable, such that two categorically similar objects were perceptually more difficult to distinguish than two categorically dissimilar objects, leading to slower visual search. To rule out this explanation, in Experiment 2 we used a same-different task to objectively measure perceptual discriminability for each pair (Jacob & Arun, 2020).

## Experiment 2

The goal of Experiment 2 was to measure the perceptual discriminability of each stimulus pair, so that we could use these values to reanalyze Experiment 1 data, regressing out a continuous measure of perceptual similarity. Participants in Experiment 2 viewed the same stimulus pairs as used in Experiment 1, intermixed with “same” trials showing the same stimulus twice (Figure 1C). However, unlike Experiment 1, in this task, there was no top-down search component: participants only had to perceptual discriminate the stimuli, indicating whether the two objects were the same or different. Performance (RT) on trials in which two different objects are presented is an index of the perceptual similarity of the two objects (Jacob & Arun, 2020; Mohan & Arun, 2012): responses will be slow for perceptually similar objects (e.g., snake-rope) and fast for perceptually dissimilar objects (e.g., snake-airplane). These data allowed us to reanalyze Experiment 1 data using regression analyses, including the perceptual similarity of each pair as a continuous variable.

### Methods

#### Participants

Twenty-four healthy volunteers (16 females, mean age = 26.75, SD = 4.16 years) recruited through social networks participated in this online experiment. This experiment was run after Experiment 3, and the sample size was matched to that of Experiment 3 (for power analysis, see Experiment 3 Participants section). All participants had normal or corrected-to-normal vision. The participants gave informed consent and received monetary compensation for their time (100 NTD, about 3 USD). The study was approved by the Ethics Committee of the Faculty of Social Sciences, Radboud University Nijmegen.

#### Stimuli

The stimuli in Experiment 2 were identical to those in Experiment 1. We used eight animate objects and eight inanimate objects. Each object had four exemplars, resulting in 64 stimuli in total. All stimuli were gray scaled and presented on a gray background. The experiment was run online and coded using the JSPsych toolbox 7.0.0 (de Leeuw, 2015).

#### Task design and procedure

The experiment consisted of two conditions: same and different objects. All analyses focused on the different object condition, consisting of the 32 pairs used in Experiment 1. In the same-object condition, the object was presented with an identical one that was 110%, 100% or 90% as large to avoid participants performing the task based on low-level cues (Arun, 2012). In the different-object condition, the two objects were both presented at their original size, similar to Experiment 1. The trial procedure followed that of Experiment 1, except for the absence of the target cue (Figure 1C). In the beginning of a trial, a 500-msec fixation cross was presented at the center of the screen to signal the onset of the trial. Then, a search display consisting of two objects was presented for 300 msec and followed by a maximal 800 msec blank for response. Participants were instructed to judge whether the two objects were the same or different. They responded by pressing keyboard buttons, pressing letter F when objects were the same and letter J when they were different. The inter-trial intervals varied randomly between 800, 1000, and 1200 msec.

#### General experimental procedure

Participants first received written instruction about the task requirements. Then, participants practiced a block of 6 trials consisting of four different objects not used for the formal experiment. During the actual experiment, participants were required to fixate on a cross at the center of the screen and respond as quickly and accurately as they could. A total of 512 trials (256 “same” trials and 256 “different” trials) were presented in 4 blocks of 128 trials. Each block consisted of 64 same-object trials and 64 different-object trials. Of the 64 same-object trials, 32 trials consisted of two objects at 100% of their original size, 16 trials consisted of one object at 100% and one object at 110% of their original size, and 16 trials consisted of one object at 100% and one object at 90% of their original size. The 64 different-object trials consisted of all 32 pairs present twice, with each object presented on each side once. Each block used one of the four exemplars, so the frequency of the presentation of exemplars was equal. Trial order within blocks was randomized. Participants could take a rest between blocks and they could self-initiate the next block. The full experimental time for each participant was between 20 to 30 minutes.

#### Behavioral analyses

We checked the accuracy data first to make sure that participants fully understood and followed the experiment instruction. We computed mean accuracy to test for outliers (accuracy lower than Mean – 3*SD); no participants had to be excluded. The analyses focused on Reaction Times (RTs) data for correct trials (Jacob & Arun, 2020). We calculated the mean RT for each stimulus pair from the different-object conditions for further trial-based model analyses. Furthermore, a paired-samples *t*-test was conducted for different-object conditions to test the categorical effect by comparing mean RT to stimulus pairs from the same category (e.g., both animate) with mean RT to stimulus pairs from different categories (animate-inanimate).

#### Model-based analyses of Experiment 1 data

We used the pairwise RT data of Experiment 2 as a model of perceptual similarity for Experiment 1. This allowed us to more conclusively dissociate categorical from perceptual similarity effects on visual search. Categorical similarity was binary, with objects either belonging to the same or different categories. Perceptual similarity consisted of the RTs from the same-different task of Experiment 2 (mean across participants of Experiment 2). The similarities were *z* transformed, and the interaction was the product of *z*-transformed categorical similarity and *z*-transformed perceptual similarity. For each participant of Experiment 1, multiple linear regression was used to predict *z*-transformed RTs using three predictors (categorical and perceptual similarity, and their interaction), thus yielding three beta estimates per participant. At the group level, one-sample *t*-tests were used to compare the beta estimates against zero (*p* <.05, one-tailed). The procedure of the analysis is illustrated in Figure 3A.

**Figure 3.**
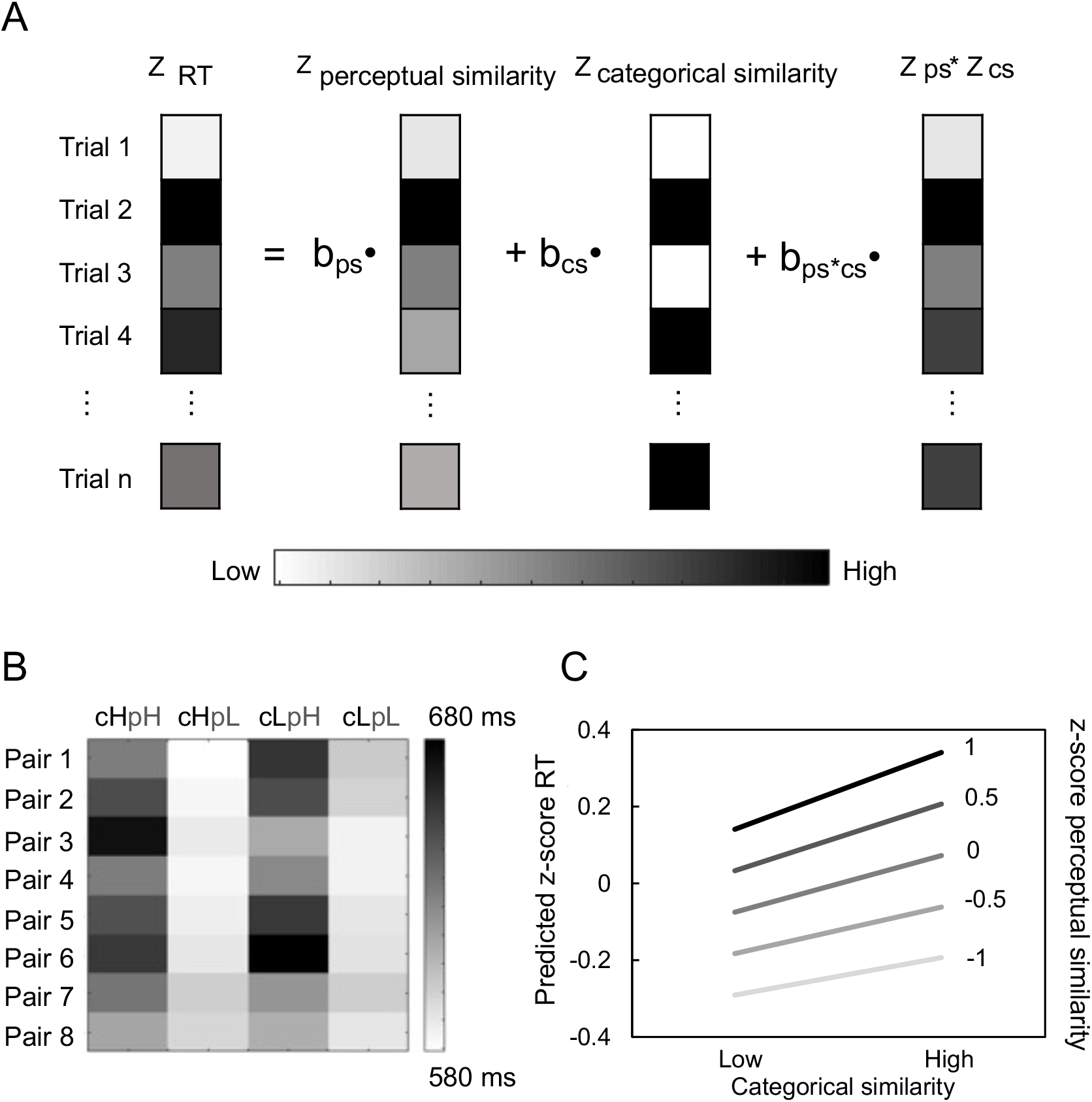
(A) Illustration of trial-based regression analysis. b_ps_ represents the slope of the perceptual similarity effect, b_cs_ represents the slope of the categorical similarity effect, and b_ps*cs_ represents the slope of the interaction between perceptual and categorical similarity effects. (B) The matrix shows the mean RTs from the same-different task (Experiment 2) for each pair of stimuli. The arrangement of pairs is identical to the matrix shown in Figure 1A. (c: categorical similarity, p: perceptual similarity, H: high similarity, L: low similarity. E.g., cHpH indicates high categorical and high perceptual similarity). (C) The predicted results are based on the beta estimates from the regression analysis. The slope is steeper when perceptual similarity increases.

### Results

#### Behavioral results

Mean accuracy was 92.58 % (*SD* = 4.10 %) and no further analysis for accuracy was conducted. Only correct trials from different-object conditions were included for RT analyses. The RT matrix for 32 pairs is illustrated in Figure 3B. No significant categorical effect was found (*t*(23) = 0.148, *p* = .442), indicating that same-category and different-category pairs were equally discriminable.

#### Trial-based regression analysis results

We conducted a trial-based regression analysis of Experiment 1 data using a continuous perceptual similarity variable established from the same-different task of Experiment 2 (Figure 3A). The results showed significantly positive beta estimates of both categorical similarity (*β* = .074, *t*(15) =8.938, *p* <.001) and perceptual similarity (*β* = 0.242, *t*(15) =11.184, *p* <.001), indicating that the higher the similarity, the slower the visual search (Figure 3C). The beta estimate of the interaction was also significant (*β* = .027, *t*(15) =3.063, *p* =.008), indicating that the categorical similarity effect was stronger when perceptual similarity increased (Figure 3C).

### Discussion

In Experiment 2, we measured perceptual similarity using a same-different task that resembled the visual search task except for the absence of a top-down search component. Data from this experiment was used to reanalyze Experiment 1 data, now including a continuous measure of perceptual similarity. The results replicated the 2-by-2 repeated-measures ANOVA of Experiment 1, again showing that categorical and perceptual similarity interact. Because perceptual similarity was directly included in the model, it is unlikely that the categorical similarity effect reflected differences in perceptual discriminability between same-and different-category pairs.

## Experiment 3

In Experiments 1 and 2, we found that both perceptual and categorical similarity influenced search performance, observing slower search when target-distractor similarity was high. Moreover, the influence of categorical similarity was modulated by perceptual similarity, being stronger when perceptual similarity was high. However, these behavioral results do not provide information about the time course of perceptual and categorical similarity effects (and their interaction). Furthermore, the behavioral results do not distinguish between influences on attentional allocation versus influences on later decision-making stages.

Therefore, in Experiment 3, we measured the target-related N2pc, a lateralized ERP component, during the visual search task of Experiment 1. The N2pc component is considered an index of spatial selective attention (Eimer, 1996; Eimer & Grubert, 2014; Mazza et al., 2009; Mazza et al., 2007; Nako et al., 2016). This component usually emerges around 200 to 300 msec after stimulus onset with more negative amplitude over posterior electrodes contralateral to the target (Eimer, 1996; Grubert & Eimer, 2016; Kiss et al., 2008; Luck & Hillyard, 1994; Mazza et al., 2007; Woodman & Luck, 1999).

According to the hierarchical processing hypothesis outlined in the Introduction, we would expect perceptual similarity effects to precede categorical effects. Previous studies showed that such differences may arise within the N2pc window, with the N2pc to a categorical target delayed by about 40 msec relative to the N2pc triggered by a specific physical target (Nako, Wu, Smith, et al., 2014) or singleton color (Callahan-Flintoft, & Wyble, 2017). Finally, mirroring the behavioral results, we expect an interaction between categorical and perceptual similarity, with stronger categorical effects when the distractor is perceptually similar to the target.

### Methods

#### Participants

Twenty-four healthy volunteers (13 females, mean age = 23.63, SD = 3.08 years) participated in Experiment 3. This sample size was required to achieve 0.8 power to detect the interaction effect of Experiment 1 (*η^2^_p_* = 0.282, Cohen’s *d* =0.607; calculated by PANGEA v0.2; Westfall, 2016). All participants had normal or corrected-to-normal vision and were right-handed according to the Edinburgh Handedness Inventory (Oldfield, 1971). The participants gave informed written consent, and received monetary compensation for their time (750 NTD, about 27 USD). The study was approved by the research ethics office of the National Taiwan University.

#### Stimuli

The stimuli and task design in Experiment 3 were identical to those in Experiment 1. Eight animate objects and eight inanimate objects were used. Each object had four exemplars, resulting in 64 stimuli in total. All stimuli were gray scaled and presented on a gray background. Each stimulus was fitted into an invisible frame subtending a visual angle of approximately 8° (height) × 8° (width). Stimuli were controlled by the E-Prime software (Psychology Software Tools, Inc.) and presented on a 17-in. cathode ray tube monitor with a refresh rate of 60 Hz.

#### Task design and procedure

The task design and procedure were also identical to those of Experiment 1 (Figure 1C). The experimental design followed a 2 (categorical similarity: high, low) × 2 (perceptual similarity: high, low) within-subjects factorial design. Each trial began with a 500-msec fixation period. Next, a target cue in Chinese was presented for 1000 msec and followed by another fixation cross for 500 msec. After that, a search display was presented for 300 msec and followed by a maximal 800 msec blank for response. The inter-trial intervals varied randomly between 800, 1000, and 1200 msec.

#### General experimental procedure

In Experiment 3, EEG data were recorded while participants performed the experiment. Participants sat in a dimly illuminated room at a distance of 57 cm from the monitor. After that, participants received instructions about task requirements and practiced a block of 8 trials. During the actual experiment, participants were required to fixate on a cross at the center of the screen and respond as quickly and accurately as they could. Participants were instructed to minimize eye blinking and movement during the task. All trial types were randomized and equiprobable within 16 blocks of 64 trials, yielding a total of 1024 trials (256 trials for each condition). The combination and counterbalance were identical to Experiment 1, but two blocks were combined into one block here. Participants could take a rest between blocks and they could self-initiate the next block. The full experimental time for each participant was between 60 to 80 minutes.

#### EEG data acquisition

Electrophysiological data were recorded using a NuAmp amplifier (Neuroscan Inc.) with 37 Ag/AgCl electrodes. The electrodes included six sites in the central line (Fz, FCz, Cz, CPz, Pz, and Oz) and 12 sites over the left and right hemispheres (FP1/FP2, F3/F4, F7/F8, FC3/FC4, FT7/FT8, C3/C4, T3/T4, CP3/CP4, TP7/TP8, P3/P4, T5/T6, and O1/O2). The vertical and horizontal eye movements were also recorded using EOG electrodes. Three additional electrodes served as the ground (AFz placed between FPz and Fz) and reference (A1 and A2 placed on each mastoid site). Impedances for each electrode was kept below 5 KΩ. Brain activity was recorded continuously with 1000-Hz analogue-to-digital sampling rate and filtered with a low-pass filter of 300 Hz. Trial-type codes were sent to the computer used to recorded EEG data via a parallel port from the computer used to present task stimuli.

#### EEG data preprocessing

Offline EEG data were preprocessed using SPM12 software (Wellcome Trust Centre for Neuroimaging, University College London, UK) and the Fieldtrip toolbox (Oostenveld et al., 2011) in MATLAB (MathWorks). EEG data were re-referenced using the average of the Al and A2 electrodes and then epoched from −500 to 1000 msec relative to stimulus onset. Epochs were baseline-corrected using the prestimulus period (from −100 to 0 msec). Epochs containing artifacts (i.e., blinks or eye-movement) were rejected based on visual inspection. Epochs with incorrect responses were also excluded in further analyses.

#### Behavioral analyses

The behavioral analyses included two-way repeated-measures ANOVA and trial-based regression analyses. Accuracy and Reaction Times (RTs) were included, but only correct trials were included in the RT analyses.

#### ERP analyses

We first tested the N2pc effect across all conditions using a paired-samples *t*-test. Following previous work (Yeh et al., 2019), we compared the difference between the mean amplitude of posterior electrodes (using the average from electrodes T5/6, TP7/8, and P3/4; note that PO7/8 do not exist in this EEG system) contralateral vs ipsilateral to the target. This analysis was restricted to 200 to 299 msec after stimulus onset based on previous research on the N2pc (Grubert & Eimer, 2016; Luck & Hillyard, 1994; Mazza et al., 2007; Woodman & Luck, 1999). In subsequent analyses, we used the N2pc effect (ipsilateral - contralateral) to test for the influence of perceptual and categorical similarity. Note that more positive values indicate a stronger (more negative) N2pc. To test our hypothesis related to early vs late influences of perceptual and categorical similarity, we divided the N2pc time window into early (200-249 msec) and late (250-299 msec) parts (for a similar approach, see Eimer & Kiss, 2008; Feldmann-Wustefeld et al., 2015; Yeh et al., 2019). The resulting data were analyzed using a three-way repeated-measures ANOVA with factors: time window (early, late), perceptual similarity (high and low), and categorical similarity (high and low).

Furthermore, to test for similarity effects with high temporal resolution, we also tested similarity effects from 200 to 299 msec with 1-ms resolution. To account for multiple comparisons, main effects and interactions were statistically tested using cluster-based nonparametric permutation tests (Maris & Oostenveld, 2007), thresholded at *p* < .05, one-tailed.

#### Trial-based model analyses for N2pc

Additionally, we tested the main effects of perceptual and categorical similarity, and their interaction, using model-based analyses on trial-by-trial basis, similar to Experiment 2. First, we calculated the N2pc by subtracting the amplitude at the contralateral side from the ipsilateral side relative to the target location for each trial for early and late time windows. Then, we fit a regression model to the *z*-transformed N2pc using *z*-transformed perceptual and categorical similarity, and their interaction at the individual-participant level. Finally, we tested beta estimates against zero using one-sample *t*-tests at the group level.

Furthermore, to increase the temporal resolution for the similarity effects, we calculated N2pc and ran the regression analysis for every 10-msec bin from 200 to 299 msec. To account for multiple comparisons, we used a cluster-based nonparametric permutation one-sample *t* tests (Maris & Oostenveld, 2007) to compare the beta estimates against zero with null distribution created from 1000 Monte Carlo random partitions and cluster correction type I error (*p* <.05, one-tailed).

### Results

#### Behavioral results

The behavioral results of Experiment 3 replicated the results of Experiment 1. The RT data (Figure 4A) revealed significant main effects of both categorical similarity, *F*(1, 23) = 109.791, *p* < .001, *η^2^_p_* = 0.827 and perceptual similarity, *F*(1, 23) = 127.471, *p* < .001, *η^2^_p_* = 0.847. As shown in Figure 4A, responses in the low categorical similarity condition (*M* = 501.742 msec, *SE* = 15.138 msec) were faster than in the high categorical similarity condition (*M* = 517.801 msec, *SE* = 15.469 msec), and the responses in the low perceptual similarity condition (*M* = 489.599 msec, *SE* = 14.666 msec) were faster than in the high perceptual similarity condition (*M* = 529.944 msec, *SE* = 16.080 msec). We again found a significant two-way interaction between categorical and perceptual similarity, *F*(1, 23) = 6.970, *p* = .015, *η^2^_p_* =0.233. To follow up on the interaction, we tested the categorical effect separately for low and high perceptual similarity conditions (Figure 4B) using paired sample *t*-tests (with Bonferroni correction for two tests, *p*<.025). The categorical effect was observed in both the high perceptual similarity condition (*t*(23) = 8.513, *p*< .001, *M*_difference_ = 21.240 msec, *SE* = 2.495 msec) and the low perceptual similarity condition *t*(23) = 4.37, *p* < .001, *M*_difference_ = 10.880 msec, *SE* = 2.485 msec).

**Figure 4.**
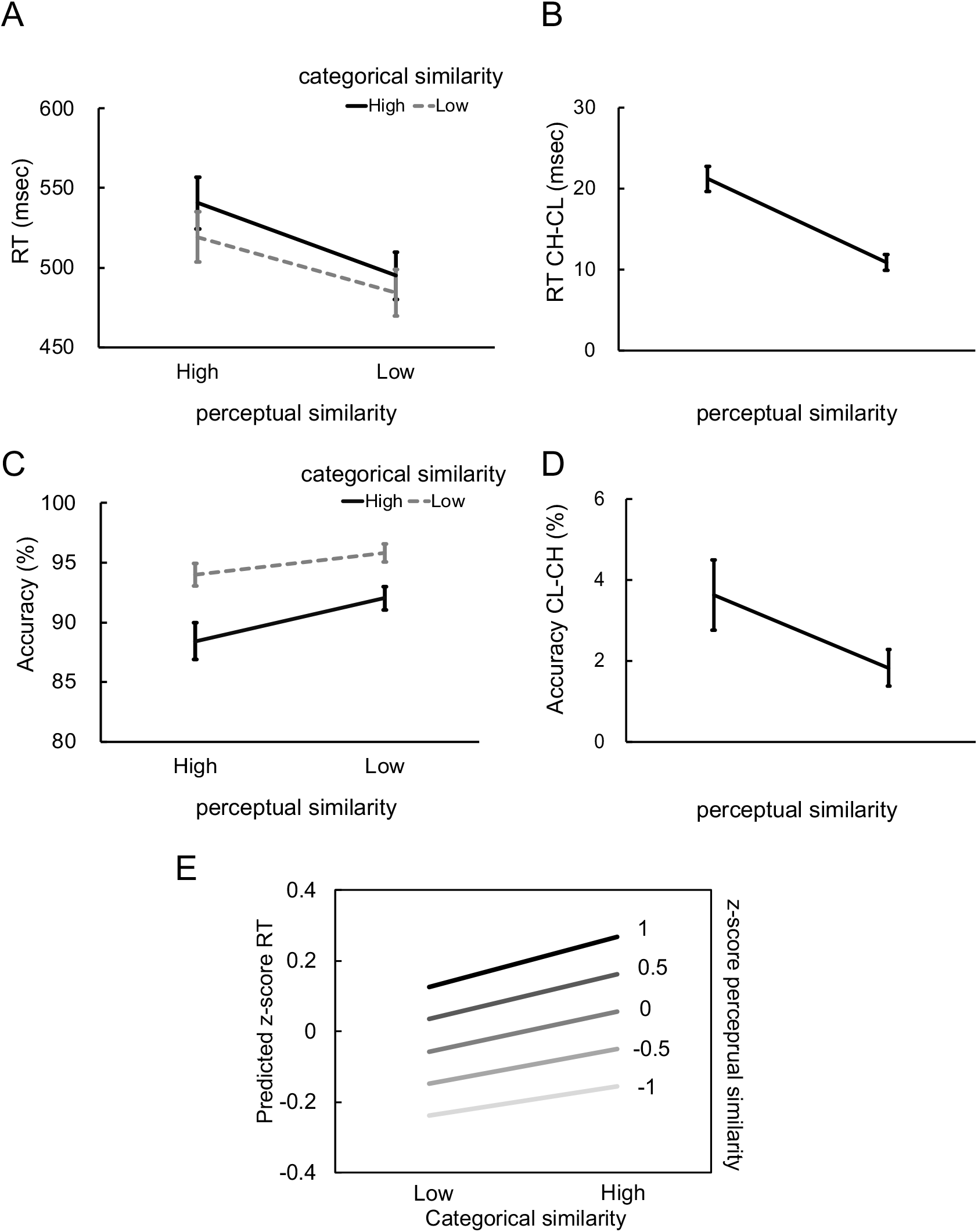
Behavioral results of Experiment 3, including mean RT (msec) in the top panel and accuracy (%) in the bottom panel. Error bars represent the standard error of the mean. (A) Reaction time results for all four conditions. (B) The categorical effect (low-high categorical similarity) of reaction time separately for high and low perceptual similarity conditions. (C) Accuracy results for all four conditions. (D) The categorical effect (low-high categorical similarity) of accuracy separately for high and low perceptual similarity conditions. (E) Predicted results based on the beta estimates from the regression analysis.

The accuracy data (Figure 4C) showed significant main effects of both categorical similarity, *F*(1, 23) = 20.008, *p* < .001, *η^2^_p_* = 0.465, and perceptual similarity, *F*(1, 23) = 45.443, *p* < .001, *η^2^_p_* = 0.664. The results indicated that the low categorical similarity condition (*M* = 93.938%, *SE* = 0.810%) was more accurate than the high categorical similarity condition (*M* = 91.208%, *SE* = 1.198%), and the low perceptual similarity condition (*M* = 94.917%, *SE* = 0.828%) was more accurate than the high perceptual similarity condition (*M* = 90.229%, *SE* = 1.209%). The interaction between categorical and perceptual similarity was also significant, *F*(1, 23) = 7.384, *p* = .015, *η^2^_p_* =0.243. To follow up on the interaction, we tested the categorical effect separately for low and high perceptual similarity conditions (Figure 4D) using paired sample *t*-tests (with Bonferroni correction for two tests, *p*<.025). The categorical similarity effect was significant when perceptual similarity was high (*t*(23) = 4.178, *p* < .001, *M*_difference_ = 3.625 %, *SE* = 0.867%) and when it was low (*t*(23) = 4.011, *p* < .001, *M*_difference_ = 1.833%, *SE* = 0.457%).

Next, the model-based analyses (illustrated in Figure 3A) showed significantly positive beta estimates for categorical similarity (*β* = .056, *t*(23) = 10.495, *p* <.001) and perceptual similarity (*β* = .197, *t*(23) = 15.873, *p* <.001), indicating that the higher the similarity, the slower the response. We also found a significantly positive beta estimate for the interaction between perceptual and categorical similarity (*β* = .015, *t*(23) = 2.163, *p* <.041), indicating that the higher perceptual similarity, the stronger the categorical effect (Figure 4E).

These results provide a replication of Experiment 1 and the model-based analysis of Experiment 2. Next, we turned to the EEG data to investigate the time course of perceptual and categorical influences on visual search.

#### ERP results

Across conditions, we found a significant N2pc effect (average of 200 to 299 msec after stimulus onset), with the amplitude over posterior electrodes ipsilateral to the target being more positive than the amplitude over posterior electrodes contralateral to the target (*t*(23) = 10.486, *p* <.001; Figure 5A). A three-way repeated-measures ANOVA on the N2pc (Figure 5B) showed significant main effects of time window (early [200-249msec], late [250-299msec]), *F*(1, 23) = 43.729, *p* < .001, *η^2^_p_* = 0.655, and perceptual similarity, *F*(1, 23) = 19.569, *p* < .001, *η^2^_p_* = 0.460. The main effect of categorical similarity was not significant, *F*(1, 23) = 3.442, *p* = .076, *η^2^_p_* = 0.13. Importantly, the results also showed a significant three-way interaction between time window, perceptual similarity, and categorical similarity, *F*(1, 23) = 11.625, *p* = .002, *η^2^_p_* = 0.336. To clarify the three-way interaction, we ran two-way repeated-measures ANOVAs for the two time windows separately (Figure 5C).

**Figure 5.**
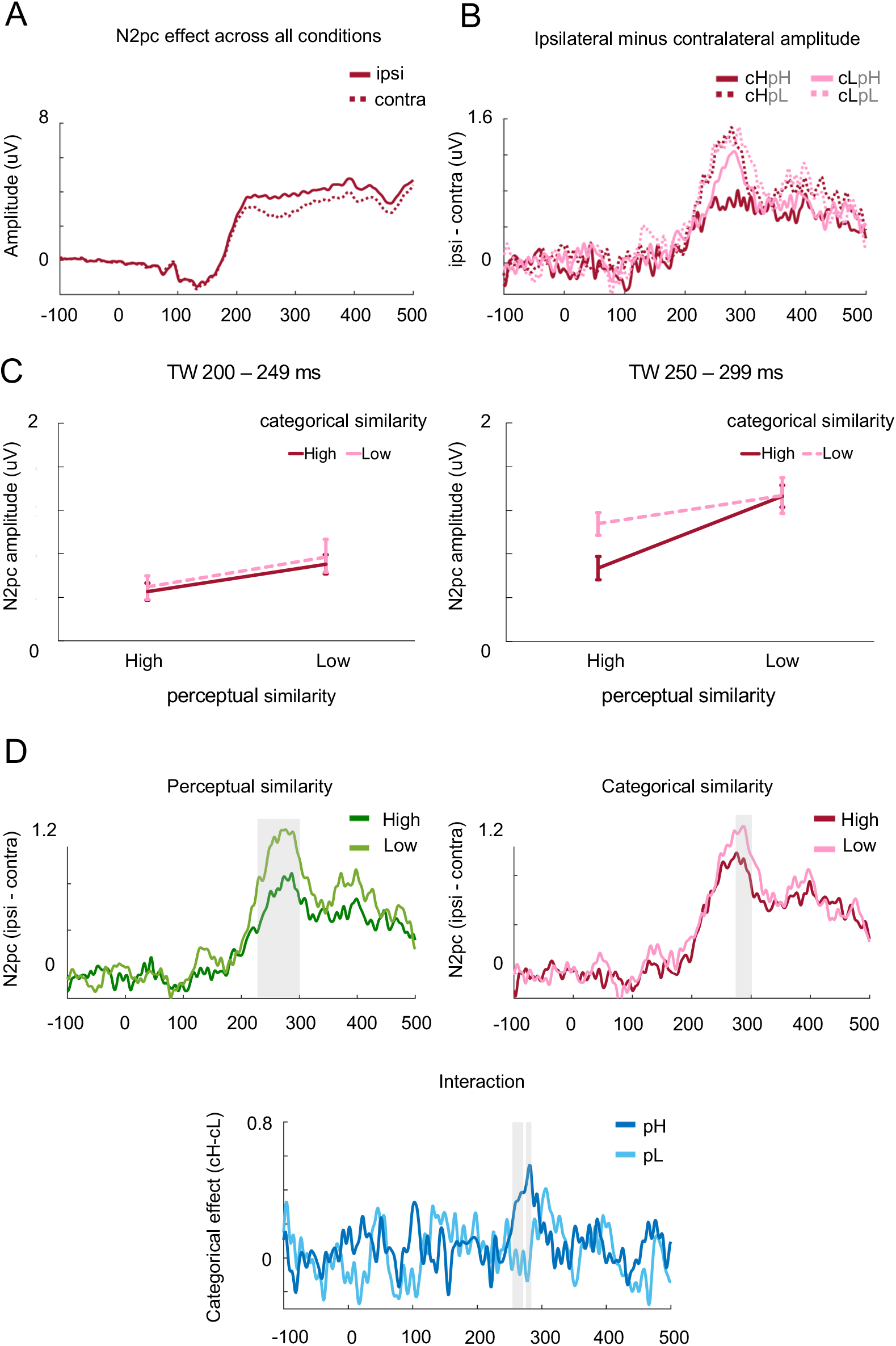
The results of ERP analyses. (A) ERP waveforms averaged across all conditions and all participants over posterior electrodes (averaging electrodes T5/6, TP7/8, and P3/4; ipsilateral waveforms: solid lines; contralateral waveforms: dashed lines). (B) The amplitude of the difference between ipsilateral and contralateral sides for the four conditions (c: categorical similarity, p: perceptual similarity, H: high similarity, L: low similarity. E.g., cHpH indicates high categorical and high perceptual similarity). (C) The N2pc effects for 2 by 2 conditions for two time windows (200-249ms; 250-299ms). (D) The results from permutation tests. Grey backgrounds indicate significant clusters. Top-left panel shows the N2pc effect (subtracting amplitude at contralateral side from ipsilateral side) for high (green) and low (light green) perceptual similarity. Top-right panel shows the N2pc effect for high (red) and low (pink) categorical similarity. Bottom panel shows the categorical effect under high (blue) and low (light blue) perceptual similarity.

For the early time window (200-249 msec), there was a significant perceptual similarity effect, *F*(1, 23) = 7.582, *p* = .011, *η^2^_p_* =0.248, but no main effect of categorical similarity, *F*(1, 23) = 0.391, *p* = .538, *η^2^_p_* =0.017, and no interaction between perceptual and categorical similarity, *F*(1, 23) = 0.024, *p* = .855, *η^2^_p_* =0.001. By contrast, for the late time window (250-299 msec), there were significant main effects of both categorical, *F*(1, 23) = 8.154, *p* = .009, *η^2^_p_* =0.262, and perceptual similarity, *F*(1, 23) = 24.556, *p* < .001, *η^2^_p_* =0.516. There was also a marginally significant two-way interaction between categorical and perceptual similarity, *F*(1, 23) = 3.94, *p* = .059, *η^2^_p_* =0.146. To follow up on the interaction, we tested the categorical effect separately for low and high perceptual similarity conditions using paired sample *t*-tests (with Bonferroni correction for two tests, *p*<.025). The results showed that the categorical similarity effect was only significant when perceptual similarity was high, *t*(23) = 3.792 *p* < .001, *M*_difference_ = 0.406, *SE* = 0.107. No significant categorical similarity effect was found when perceptual similarity was low *t*(23) = 0.055 *p* = .478, *M*_difference_ = 0.008, *SE* = 0.138. The ANOVA results of ERP analyses are illustrated in Figure 5C.

Using cluster-based nonparametric permutation tests (constrained to the N2pc window, 200-299 msec), we found a significant perceptual similarity effect from 224 to 299 msec (cluster-based *p* <.001) and a significant categorical similarity effect from 280 to 299 msec (cluster-based *p* <.001). We also found two significant clusters for the interaction, from 250 to 268 (cluster-based *p* =.024), and from 274 to 282 (cluster-based *p* =.025). The permutation test results are illustrated in Figure 5D.

#### Trial-based model analyses results for N2pc

In trial-based model analysis, using the RT data of Experiment 2 as a continuous measure of perceptual similarity, in the early time window (200-249 msec), we found a significantly negative beta estimate for the perceptual similarity predictor (*β* = −.024, *t*(23) = −3.368, *p* =.001), indicating that increasing perceptual similarity reduced the N2pc. The main effect of categorical similarity (*β* = −.002, *t*(23) = −0.275, *p* =.393) and the interaction between categorical and perceptual similarity (*β* = −.002, *t*(23) = −0.403, *p* =.345) were not significant. In the late time window (250-299 msec), both main effect betas were significantly different from zero: perceptual (*β* = −.038, *t*(23) = −5.106, *p* <.001), categorical (*β* = −.013, *t*(23) = −2.361, *p* =.014). Most importantly, there was a significant interaction between perceptual and categorical similarity (*β* = −.020, *t*(23) = −2.603, *p* =.007), indicating that the higher the perceptual similarity the more negative was the beta estimate for the categorical similarity predictor (Figure 6A).

**Figure 6.**
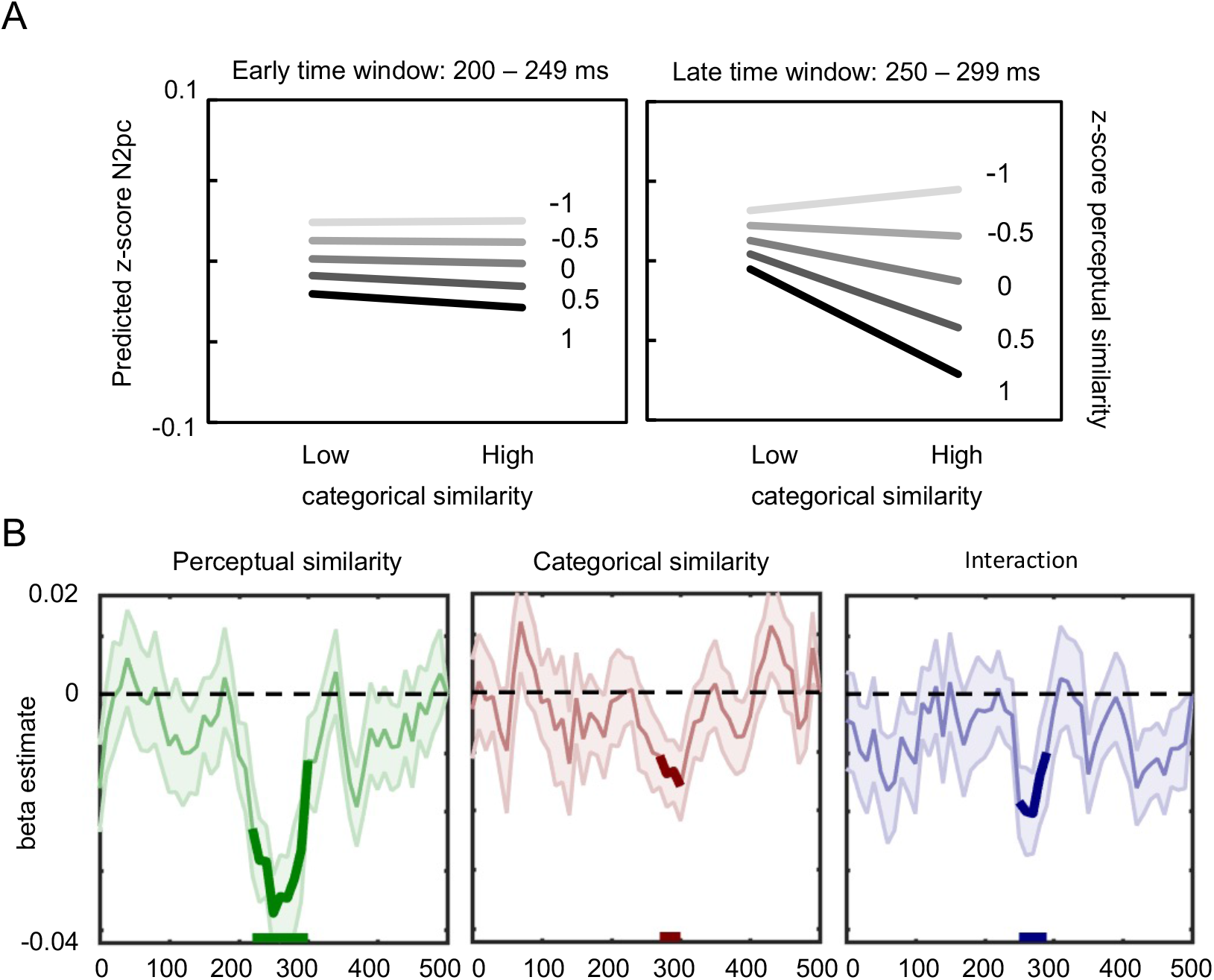
(A) Predicted results based on the beta estimates from the regression analysis. The left panel shows the results for time window 200 to 249 msec. The right panel shows the results for time window 250 to 299 msec. (B) Significant results from cluster-based permutation tests. The bold lines represent the significant clusters (cluster-based *p*-value <.05, one-tailed).

Finally, in clustered-based permutation tests (constrained to the N2pc time window, 200-299 msec), we identified a main effect of perceptual similarity from 210 to 299 msec (cluster-based *p* <.001) and a main effect of categorical similarity from 270 to 299 msec (cluster-based *p* =.031). These effects indicated that the lower the similarity the stronger the N2pc. The interaction between perceptual and categorical similarity was significant from 250 to 289 msec (cluster-based *p* =.025). The interaction indicates that the categorical similarity effect was stronger when perceptual similarity was high. The results are illustrated in Figure 6B.

### Discussion

In Experiment 3, we used EEG to measure the time course of influences from perceptual and categorical target-distractor similarity during visual search. Mirroring the behavioral results, we found that the N2pc (an index of attentional allocation) was influenced both by perceptual and categorical similarity. Categorical similarity again interacted with perceptual similarity, with stronger categorical similarity effects when perceptual similarity was high. Furthermore, the ERP results showed a clear distinction in the time-course of perceptual and categorical influences. While the influence of perceptual similarity started as early as 210/220 ms after stimulus onset, the categorical effect emerged only at around 270/280 ms. Results of the trial-based regression analysis (Figure 6) indicated that the late categorical similarity effect could not be attributed to residual perceptual similarity differences between conditions.

Importantly, this categorical effect was qualified by an interaction with perceptual similarity, being observed only when perceptual similarity was high. These results are in line with our hierarchical processing hypothesis, with attention initially guided by perceptual stimulus attributes, followed by categorical attributes.

## General Discussion

The current behavioral and electrophysiological results demonstrate how the search for a target object is influenced by both the perceptual and the categorical similarity of a concurrently presented distractor object. Behaviorally, search was slowest when the distractor was both perceptually and categorically similar to the target. An interaction between perceptual and categorical similarity revealed that the influence of categorical similarity was strongest when the distractor was perceptually similar to the target. The same interaction was also observed in the late part of the N2pc (250-300 ms after onset), an ERP component indexing spatial attention. Here, we observed that perceptual and categorical similarity influenced spatial attention sequentially: perceptual similarity affected spatial attention before categorical similarity did. Furthermore, the categorical effect was qualified by an interaction with perceptual similarity, being observed only for perceptually similar distractors. Thus, by orthogonally manipulating perceptual and categorical target-distractor similarity, we demonstrate 1) that purely categorical similarity affects spatial attention during visual search, and 2) that such categorical influences follow perceptual influences, being primarily observed when perceptual properties are ineffective in guiding attention to the target.

The focus on categorical target-distractor similarity effects during visual search follows a long line of research testing for such effects. However, many of these studies could not clearly distinguish between categorical and perceptual similarity. For example, the classical alphanumeric category effect (Jonides and Gleitman, 1972), in which search for a letter (O) or a digit (0) is more efficient in arrays consisting of items of the opposite category proved difficult to replicate, and may have reflected physical/perceptual similarity rather than categorical similarity (Krueger, 1984; Cardosi, 1986). As reviewed in the Introduction, studies investigating visual search for objects have also provided evidence for categorical and semantic effects in visual search (Belke et al., 2008; Moores et al., 2003; Telling et al., 2010). However, also in these studies, it was hard to rule out residual perceptual influences. A previous study that, similar to the current study, investigated search for animate and inanimate objects (Levin et al., 2001) showed that the between-category advantage (e.g., search for an animal among objects) could be explained by target-distractor differences in global contour shape (e.g., rectilinearity). In the present study, to control for such perceptual differences, we included a same-different task to objectively measure the perceptual similarity of each target-distractor pair (Jacob & Arun, 2020). This task was identical to the visual search task, except that it lacked a top-down search component. Similarity between objects in terms of their perceptual properties (e.g., shape) influences performance in the same-different task; e.g., participants are slower to respond “different” to two objects with a similar shape than to two objects with a dissimilar shape. Importantly, categorical similarity effects were observed in the visual search task even when regressing out this measure of perceptual similarity. Therefore, the categorical similarity effects observed in the current study are unlikely to reflect perceptual similarity effects.

The key finding of the current study was that the influence of categorical similarity was primarily observed when targets and distractors were perceptually similar. This interaction was most clearly observed in the N2pc indexing spatial attention; here, the categorical similarity effect was exclusively observed when perceptual similarity was high (e.g., Figure 5C). Unlike the N2pc results, the behavioral results also showed a (reduced) categorical similarity effect when perceptual similarity was low. This suggests that categorical similarity may influence search beyond the attentional selection stage (as indexed by the N2pc). For example, categorical similarity may additionally affect identification and decision-making stages of the search process (Eimer, 2014; Wolfe, 2021).

The interaction observed in the N2pc can be interpreted in at least two ways. First, it is possible that category guides search when search cannot be guided by perceptual properties. On this *category-before-attention* account, when two perceptually similar objects are presented, neither object is selected and both objects are processed to the category level (within 250 ms). When category is informative, i.e., when the target is of a different category than the distractor, attention is then guided by category, giving rise to a lateralized EEG response over posterior electrodes. This account is consistent with prior work that proposed that category can guide attention (Nako et al., 2016; Nako, Wu, & Eimer, 2014; Nako, Wu, Smith, et al., 2014). Different from these previous studies, however, here, participants were not instructed to search for targets at the (super-ordinate) category level, but instead searched for basic-level categories (e.g., snake). Super-ordinate (animate-inanimate) categorical similarity effects were observed even though the same-category distractor item was always from a different basic-level category (e.g., snail). This suggests that the target’s super-ordinate category was automatically included in the search template.

An alternative account of the current N2pc findings is that it reflects differential disengagement of attention as a function of categorical similarity. On this *attention-before-category* account, spatial attention is not guided by category. When two perceptually similar objects are presented, perceptual guidance fails to direct attention to the target, such that attention is randomly directed to either the target or the distractor object, irrespective of categorical similarity. On trials where the distractor object is selected, attention may be slower to disengage from this object when its category matches that of the target than when it does not, for example, because it better matches a categorical target template held in activated long-term memory (Wolfe, 2021). This lingering of spatial attention at the distractor location could then express as reduction of the late part of the target-related N2pc.

One way in which future work could distinguish between these accounts is by increasing the competition between items, e.g., by increasing the number of distractors. In the current study, only two objects were presented simultaneously (one in each hemifield), such that both objects may have been processed to the category level even without attentional selection. When the number of items in the display increases, however, competition increases and higher-level object properties may no longer be extracted for all objects (Desimone & Duncan, 1995). In this case, (low-level) perceptual properties will remain effective in guiding attention, as extracting these does not require spatial attention (Treisman & Gelade, 1980). However, when one of the distractors is perceptually similar to the target, categorical information may now not be available to guide attention to the target. Accordingly, the first account predicts that there would be no category-related N2pc effect; attention would move serially to the perceptually-similar items without a categorical influence. By contrast, the second account would predict a qualitatively similar pattern of results as observed here: attention would still be slower to disengage from a categorically-similar than a categorically-dissimilar distractor.

In the current study, we used animacy as the categorical variable to maximize our chances of finding a categorical influence on visual search: the animate-inanimate distinction is one of the main organizing principles in visual cortex (Chao et al., 1999; Downing et al., 2006; Kriegeskorte et al., 2008) and the semantic system (Warrington & Shallice, 1984; Caramazza & Shelton, 1998). Furthermore, a previous fMRI study demonstrated that animacy is represented independently of perceptual similarity in visual cortex (Proklova et al., 2016). Future studies will need to test whether the categorical similarity effects observed here are specific to animacy and/or to other categories that are selectively represented in the visual cortex (e.g., faces, bodies, words, tools; Downing et al., 2006). It is plausible that other category distinctions (e.g., vehicles vs furniture) may not be processed rapidly and automatically enough to guide attention.

In conclusion, the current study demonstrated how low-level perceptual and high-level categorical attributes together influence visual search. We provided evidence for hierarchical processing in visual search: spatial attention was first guided by perceptual properties and was subsequently influenced by categorical information – but only when there were no perceptual properties to guide attention.

## Acknowledgements

This work was supported by a grant from the Ministry of Science and Technology, Taiwan (MOST 109-2917-I-002-023) and the European Research Council under the European Union’s Horizon 2020 research and innovation program (Grant Agreement No. 725970). The authors thank Dr. Bo-Cheng Kuo for providing EEG equipment to collect data, Dr. Yei-Yu Yeh for giving advice on the original research idea, and members of the Peelen Lab for discussion and feedback during lab meetings.

## Data availability

All stimuli and data can be found on: https://osf.io/uzp9e/

